# Chromosome-level genome assembly of macroalgae *Gracilariopsis lemaneiformis*

**DOI:** 10.64898/2026.04.28.721235

**Authors:** Yiyi Hu, Youtao Huang, Yushun Yong, Erlei Shang, Bixia Zhang, Zhenghong Sui

**Affiliations:** Key Laboratory of Marine Genetics and Breeding (Ocean University of China), Ministry of Education, Qingdao 266003, China

## Abstract

As an important cultivated red alga, *Gracilariopsis lemaneiformis* has great economic and ecological value. However, its existing genome assembly is highly fragmented and inadequately annotated. In this study, we constructed the first high-quality chromosome-level genome of *Gp. lemaneiformis* using PacBio long reads, Illumina short reads and Hi-C sequencing data. The assembled genome was approximately 86.66 Mb and the assembled sequences were anchored to 28 pseudo-chromosomes with lengths ranging from 1.70 to 7.81 Mb. 99.91% of the PacBio reads could be mapped to our assembly. In total, 8,664 genes were annotated, and the repeat elements identified in *Gp. lemaneiformis* constituted 65.04% of the whole genome, including 2.24% tandem repeat sequences and 62.81% interspersed repeats. We also established a high-evidence phylogenetic tree from 19 representative algae species, with the main aim to calculate their divergence times. This high-quality genome of *Gp. lemaneiformis* provides a crucial foundation for understanding genetic characteristics, investigating the genomic evolution, and facilitating molecular breeding.

## Introduction

*Gracilariopsis lemaneiformis* is a commercially important species from the Rhodophyta. It serves as a vital source of agar, which is extensively utilized in food processing, chemical manufacturing, pharmaceutical, cosmetic and biotechnology industries^1^. *Gp. lemaneiformis* is also main feed for abalone aquaculture^2^. In addition, *Gp. lemaneiformis* can reduce the contents of nitrogen and phosphorus in the seawater, effectively alleviating eutrophication^3,4^. Currently, *Gp. lemaneiformis* is widely cultivated across Fujian, Shandong and Guangdong provinces in China with a cultivation area exceeding 10,000 hectares. The annual production (dry weight) reaches nearly 600,000 tons, making it the second largest cultivated seaweed after *Saccharina* in China^5^.

*Gp. lemaneiformis* is an ideal material for theoretical research. This species has a typical *Polysiphonia* type life history, consisting of a gametophytic phase (female and male gametophytes) and two sporophytic phases (tetrasporophyte and carposporphyte)6. Its whole life history can be completed in a laboratory culture7. As long as the culture conditions are suitable, diploid (carpospore and tetrasporophyte) and haploid (tetraspore and gametophyte) can be obtained at any time for observation, statistics, hybridization, mutagenesis, genetic recombination and progeny analysis. In recent years, intensive research has been carried out on *Gp. lemaneiformis*, including functional gene studies^8,9,10,11^, mutation research^12,13,14,15^, genetic diversity analyses16,17,18,19,20, mitochondrial genome21, chloroplast genome22 and genomic studies^23,24^.

The species of red algae whose genome have been published include *Cyanidioschyzon merolae*25, *Porphyridium purpureum*26, *Galdieria sulphuraria*27, *Chondrus crispus*28, *Porphyra umbilicalis*^29^, *Gracilaria changii*^30^, *Gracilariopsis chorda*^31^, *Pyropia haitanensis*32 and *Pyropia yezoensis*33. The genome size of red algae is smaller than that in plants, but it usually contains a large number of repetitive sequences which complicates the genome assembly process. In our previous reports, a draft genome assembly of *Gp. lemaneiformis* was generated with assistance of only Illumina short-read sequencing, which consisted of 125,685 scaffolds with an N50 length of 20.01 kb^23^. In 2018, Sun et al. constructed the *Gp. lemaneiformis* genome with the assembly of 13,825 scaffolds and N50 length of 30.59 kb24. However, the existing genome versions lag significantly behind the increasing demand for comprehensive investigation of *Gp. lemaneiformis*. Genomes characterized by a low contig N50 tend to be highly fragmented and be inadequately annotated. A high-quality genome assembly is essential for identifying functional genes and regulatory elements, as well as for investigating the genomic evolution of marine algae.

In this study, the first high-quality chromosome-level genome of *Gp. Lemaneiformis* was constructed using PacBio, Illumina and Hi-C technologies. The final assembly was 86.66 Mb in total length, which was anchored to 28 chromosomes. A total of 8,664 protein-coding genes were predicted in the genome of *Gp. lemaneiformis*. 65.04% of the genome was composed of various repetitive elements. The chromosome-level genome assembly and comprehensive characterization of *Gp. lemaneiformis* provide valuable resources for in-depth investigations into the biological aspects of growth, development, reproduction and evolution. In addition, as an important aquaculture species, a high-quality genome serves as an crucial theoretical foundation for elucidating the genetic architecture of economic traits, conducting genetic improvement operations, and ultimately achieving cultivar selection and molecular breeding.

## Methods

### Sample preparation and sequencing

The female gametophyte named ♀6 was produced from Cultivar Lulong No. 1 (tetrasporophyte) and the thallus was cultured continuously in modified Provasoli’s medium34 under the following conditions: temperature of 20°C, photon flux density of 15 μmol m^-2^s^-1^, salinity of 30‰ and a 12 h light:12 h dark photoperiod. Approximately 4.5 g thallus was used to extract genomic DNA using a Plant Genomic DNA Kit (Tiangen Biotech, Beijing, China) following the manufacturer’s instructions.

For Illumina sequencing, at least 1μg genomic DNA was used for library construction. Libraries with insert sizes of ∼350bp were prepared following Illumina’s standard genomic DNA library preparation procedure. The qualified Illumina pair-end library would be used for Illumina HiSeq 4000 sequencing (PE=150 bp). For PacBio sequencing, 8μg DNA was spun in a Covaris g-TUBE (Covaris, MA), and then DNA fragments were purified, end-repaired and ligated with SMRT bell sequencing adapters following manufacturer’s recommendations (Pacific Biosciences, CA). 20 kb insert libraries were generated and sequenced on a PacBio Sequel instrument following the manufacturer’s instructions.

To construct Hi-C library, the fresh thallus was fixed with formaldehyde and lysed, and then the cross-linked DNA was digested with MboI overnight. Sticky ends were biotinylated and proximity-ligated to form chimeric junctions and then physically sheared to a size of 500-700 bp. Chimeric fragments representing the original cross-linked long-distance physical interactions were then processed into paired-end sequencing libraries, and 150 bp paired end reads were produced on the Illumina HiSeq 6000 platform.

For genome annotation, transcriptome sequencing was performed. Total RNA from branches of the same thallus was isolated using a Plant RNA Kit following the manufacturer’s instructions. Libraries were prepared using the Illumina TruSeq RNA library Prep kit and sequenced on an Illumina HiSeq 4000.

### Genome assembly and chromosome anchoring

The raw pair-end reads were trimmed to remove the adaptors and low-quality bases using Trimmomatic35 after quality control by FastQC (https://www.bioinformatics.babraham.ac.uk/projects/fastqc). The trimmed reads were used to do further analysis. The contig-level assembly incorporated sequencing data from a mixture of sequencing technologies, including Illumina HiSeq 4000 and PacBio Sequel as well as Hi-C scaffolding. For PacBio assembly, Canu36 was used, as it is capable of avoiding collapsed repetitive regions. Self-correction was performed which allowed us to correct all of the input PacBio reads. Then we used Pilon37 to polish the assembled genome with Illumina reads. Chromosomal assembly was constructed using the program ALLHIC38. Briefly, the Chromosomal assembly consists of the following processes: 1) Mapping Hi-C reads to the draft assembly by bwa39, and filtering the BAM file which with low quality read pairs by PreprocessSAMs.pl (LACHESIS^40^). 2) Separate all the contigs into separate clusters by the partition procedure, based on the number of links between them. 3) Reconstruct the highest scoring ordering and orientations for each cluster. 4) Building the Hi-C based chromosome-level assembly, and plot the heatmap to show the chromosome contacts.

### Repeat prediction

A *de novo* repeat library of the genome was customized using RepeatModeler 1.0.11^41^, which can automatically execute two *de novo* repeats fnding programs, including RECON v1.08^42^ and RepeatScout v1.0.5^43^. The consensus transposable element (TE) sequences generated above were imported to RepeatMasker v4.0744 to identify and cluster repetitive elements. To identify tandem repeats within the genome, the Tandem Repeat Finder (TRF) package v4.0945 was used with the modifed parameters of ‘2 7 7 80 10 50 2000 -d -h’ to find high-order repeats.

### Gene predict and annotation

The repeat-masked genome sequences were used for gene prediction. The gene prediction pipeline combined *de novo* gene predictions, homologous sequence searching and transcriptome sequence mapping. The results from the three independent methods were merged into the final consensus of gene models using EVidenceModeler v1.1.1^46^. Specifically, Augustus v3.2.3^47^ was employed for *de novo* gene prediction using appropriate parameters. Homologous sequence searching was performed by comparing protein sequences of closed related algae against the repeat-masked genome sequences using TBLASTN with parameters of identity >=60 of the query sequence covered in the alignments. The alignments were further processed using GeneWise48 to extract accurate exon-intron information. Illumina RNA-seq reads were aligned to the genome sequences using BLAT^49^. Final gene sequences were derived through further analysis of the BLAT alignment results using PASA50.

Annotation of the predicted genes was performed by blasting the protein sequences against a number of databases, including Nr, Swiss-Prot, COG, GO and KEGG, using an e-value cutoff of 1e-5. Briefly, the 200 top-scoring search results of predicted proteins against Nr, Swiss-Prot, COG and KEGG were scored based on alignment scores. The highest-scoring description was assigned to each predicted gene. GO terms were assigned to the annotated genes using the Blast2GO pipeline^51^.

### Gene clusters and phylogenetic analysis

We identified gene families by performing BLASTp search against the protein sequences of 19 algae and plants with available whole-genome information. Among these algae, 6 belonged to Rhodophyta (*Gracilariopsis chorda, Chondrus crispus, Porphyra umbilicalis, Pyropia yezoensis, Cyanidioschyzon merolae, Galdieria sulphuraria*); 1 was Phaeophyta (*Ectocarpus siliculosus*); 6 were Chlorophyta (*Chlamydomonas reinhardtii, Chlorella variabilis, Volvox carteri, Coccomyxa subellipsoidea, Ostreococcus lucimarinus, Ostreococcus tauri*); 2 were Bacillariophyta (*Phaeodactylum tricornutum, Thalassiosira pseudonana*); 1 was Glaucophyta *(Cyanophora paradoxa*) and 2 were plants (*Physcomitrella patens,Arabidopsis thaliana*). To further examine the genome divergence and conservation among algae, we carried out a phylogenetic analysis based on single-copy orthologous groups using the *Gp. lemaneiformis* genome and other genomes to build orthologous genes using OrthoMCL v2.0.352. JModeltest v2.1.1053 was employed to reconstruct a maximum likelihood phylogenetic tree for each cluster with an evolutionary model specified as ‘GTR+I+G’and to perform a bootstrap significance test with 1000 replicates.

### Gene family expansion and contraction

Based on the identified orthologs and the phylogenetic tree, gene expansion and contraction analysis was carried out using CAFE (v4.2.1) following the CAFE manual45. Gene family expansion and contraction were analyzed using *Gp. lemaneiformis* and 8 other species: *Gracilariopsis chorda, Chondrus crispus, Porphyra umbilicalis, Cyanidioschyzon merolae, Galdieria sulphuraria, Ostreococcus lucimarinus, Chlamydomonas reinhardti* and *Arabidopsis thaliana*.

## Results and Discussion

### Genome sequencing and assembly

A total of 15.6 Gb of high-quality Illumina reads, 15.5 Gb of PacBio long reads and 13.6 Gb of HiC reads were generated, representing an ∼170X coverage of the *Gp. lemaneiformis* genome (Table 1). The genome sequences were assembled into 1,350 contigs, with a N50 length of 126,001 bp. Subsequently, the contigs were anchored to 28 pseudo-chromosomes (Figure 1) with lengths ranging from 1.70 to 7.81 Mb (Table 2). The total length of the genome assembly was 86.66 Mb. The average GC content of this genome was 49.12%.

**Table 1.** Summary of the *Gp. lemaneiformis* genome sequencing and assembly.

| Items | Statistical result |
| --- | --- |
| **Sequencing** |  |
| Illumina sequencing |  |
| Clean data (Gb) | 15.61 |
| Sequencing depth (X) | 180 |
| PacBio sequencing |  |
| Clean data (Gb) | 15.50 |
| Sequencing depth (X) | 179 |
| Hi-C sequencing |  |
| Clean data (Gb) | 13.64 |
| Sequencing depth (X) | 157 |
| **Assembly features** |  |
| Assembly length (Mb) | 86.66 |
| Number of Chromosome | 28 |
| Chromosome N50 (Mb) | 3.05 |
| Chromosome N90 (Mb) | 2.02 |
| Longest chromosome (Mb) | 7.81 |
| Smallest chromosome (Mb) | 1.70 |
| Number of contigs | 1,350 |
| Contig N50 (bp) | 126,001 |
| Contig N90 (bp) | 72,000 |
| Longest contig (bp) | 1,759,000 |
| Smallest contig (bp) | 1,000 |
| Gaps (bp) | 132,200 (0.1526%) |
| Anchored rate (%) | 100 |

**Table 2.** Chromosomal characteristics of *Gp. lemaneiformis*.

| Chr ID | Chr length (Mb) | GC Content % | Gene number | mRNA length (bp) | Gene length % | Transposable element (TE) number | TE length | TE length % | Tandem repeats (TR) number | TR length | TR length % |
| --- | --- | --- | --- | --- | --- | --- | --- | --- | --- | --- | --- |
| Chr1 | 7.81 | 48.83 | 438 | 512592 | 6.56 | 3729 | 5582885 | 71.46 | 785 | 91626 | 1.17 |
| Chr2 | 4.86 | 49.76 | 698 | 954534 | 19.66 | 1947 | 2615743 | 53.87 | 426 | 48423 | 1.00 |
| Chr3 | 4.30 | 49.15 | 499 | 696342 | 16.18 | 1656 | 2503936 | 58.20 | 375 | 38785 | 0.90 |
| Chr4 | 3.86 | 48.91 | 517 | 712113 | 18.44 | 1706 | 2029942 | 52.55 | 349 | 29738 | 0.77 |
| Chr5 | 3.78 | 48.90 | 289 | 368421 | 9.74 | 1716 | 2484697 | 65.68 | 335 | 109175 | 2.89 |
| Chr6 | 3.60 | 48.94 | 226 | 269874 | 7.49 | 1609 | 2507839 | 69.59 | 313 | 51827 | 1.44 |
| Chr7 | 3.51 | 49.51 | 423 | 609261 | 17.38 | 1345 | 1960214 | 55.90 | 310 | 207285 | 5.91 |
| Chr8 | 3.27 | 48.88 | 227 | 263478 | 8.06 | 1508 | 2256197 | 69.02 | 320 | 222579 | 6.81 |
| Chr9 | 3.25 | 48.96 | 310 | 404862 | 12.46 | 1528 | 2012877 | 61.94 | 321 | 39436 | 1.21 |
| Chr10 | 3.15 | 49.42 | 390 | 536673 | 17.01 | 1191 | 1831657 | 58.06 | 259 | 26515 | 0.84 |
| Chr11 | 3.05 | 48.97 | 194 | 240249 | 7.88 | 1454 | 1963651 | 64.42 | 292 | 64982 | 2.13 |
| Chr12 | 3.01 | 48.48 | 132 | 150858 | 5.01 | 1745 | 2165993 | 72.00 | 255 | 32182 | 1.07 |
| Chr13 | 2.98 | 49.14 | 332 | 415038 | 13.91 | 1328 | 1733860 | 58.11 | 350 | 32452 | 1.09 |
| Chr14 | 2.96 | 48.54 | 220 | 269427 | 9.12 | 1464 | 1911428 | 64.67 | 294 | 48996 | 1.66 |
| Chr15 | 2.93 | 49.44 | 347 | 483222 | 16.47 | 1181 | 1676470 | 57.14 | 304 | 32208 | 1.10 |
| Chr16 | 2.91 | 49.37 | 193 | 233970 | 8.05 | 1400 | 2010959 | 69.15 | 257 | 49684 | 1.71 |
| Chr17 | 2.82 | 48.87 | 246 | 343764 | 12.17 | 1014 | 1729735 | 61.24 | 290 | 99722 | 3.53 |
| Chr18 | 2.81 | 48.82 | 143 | 157968 | 5.62 | 1295 | 1980459 | 70.46 | 237 | 26885 | 0.96 |
| Chr19 | 2.63 | 49.51 | 401 | 555999 | 21.15 | 1068 | 1200341 | 45.66 | 232 | 23428 | 0.89 |
| Chr20 | 2.61 | 49.36 | 368 | 487878 | 18.72 | 964 | 1413389 | 54.22 | 220 | 21928 | 0.84 |
| Chr21 | 2.48 | 49.53 | 278 | 359982 | 14.54 | 1041 | 1434116 | 57.94 | 216 | 23508 | 0.95 |
| Chr22 | 2.39 | 48.97 | 292 | 404832 | 16.97 | 1018 | 1265961 | 53.06 | 226 | 26460 | 1.11 |
| Chr23 | 2.25 | 49.23 | 232 | 322215 | 14.32 | 953 | 1410904 | 62.69 | 217 | 20865 | 0.93 |
| Chr24 | 2.02 | 49.77 | 230 | 353286 | 17.50 | 866 | 1114428 | 55.21 | 206 | 19767 | 0.98 |
| Chr25 | 1.99 | 49.23 | 269 | 371181 | 18.67 | 885 | 1066172 | 53.62 | 148 | 21036 | 1.06 |
| Chr26 | 1.87 | 48.90 | 138 | 196683 | 10.54 | 887 | 1227884 | 65.80 | 229 | 29078 | 1.56 |
| Chr27 | 1.86 | 49.08 | 175 | 247278 | 13.28 | 927 | 1097309 | 58.93 | 168 | 24403 | 1.31 |
| Chr28 | 1.70 | 48.90 | 85 | 105756 | 6.21 | 842 | 1227896 | 72.06 | 134 | 19697 | 1.16 |

**Figure 1.**
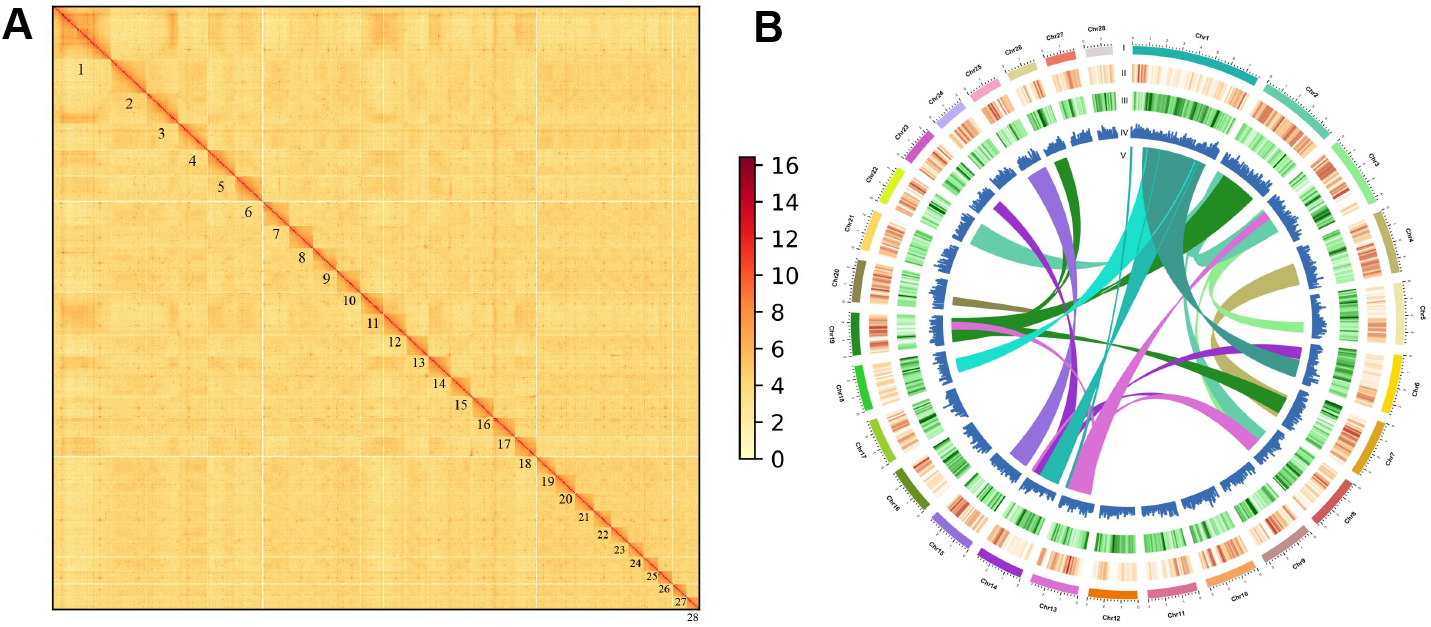
Chromatin interaction and genomic landscape of *Gp. lemaneiformis*. (A)The heatmap view of Hi-C clustering of *Gp. lemaneiformis*. (B) A circos plot of *Gp. lemaneiformis* genome. (I) Chromosome length; (II) gene density in 100 kb sliding windows; (III) repeat density in 100 kb sliding windows; (IV) GC content in 100 kb sliding windows; (V) collinear region.

The completeness of this final genome assembly was evaluated by Benchmarking Universal Single-Copy Orthologs (BUSCO), revealing that 88.4% BUSCOs are complete. For the assessment of the correctness of the genome assembly, we re-aligned PacBio reads against the assembly, then 99.91% of the reads could be mapped to the assembly sequences. Moreover, the alignment of transcriptome data against the genome assembly revealed mapping rate of 96.23%, 96.04% and 95.95%, respectively.

Compared with the assembly results of the published red macroalgae, including *Gracilariopsis chorda* (scaffold N50 = 220.2 kb), *Chondrus crispus* (scaffold N50 = 240.0 kb) and *Porphyra umbilicalis* (scaffold N50 = 202.0 kb), the assembly of *Gp. lemaneiformis* genome had the fewest contigs, the longest chromosome N50 and the highest coverage. A high-quality genome assembly is valuable for identifying structural variants, resulting in insights into the mode and tempo of genome evolution and elucidating the genetic architecture of important traits.

### Repeat prediction

For the repeat element analysis, the results showed that the repeat elements identified in *Gp. lemaneiformis* constituted 65.04% of the whole genome, including 2.24% tandem repeat sequences and 62.81% interspersed repeats (Table 3). Among the tandem repeats, 6,862 minisatellite and 1,336 microsatellites were identified. LTR elements represented the majority of the confirmed interspersed repeats, occupying 38.72% of the genome, while the DNA elements comprised 7.28%.

**Table 3.**
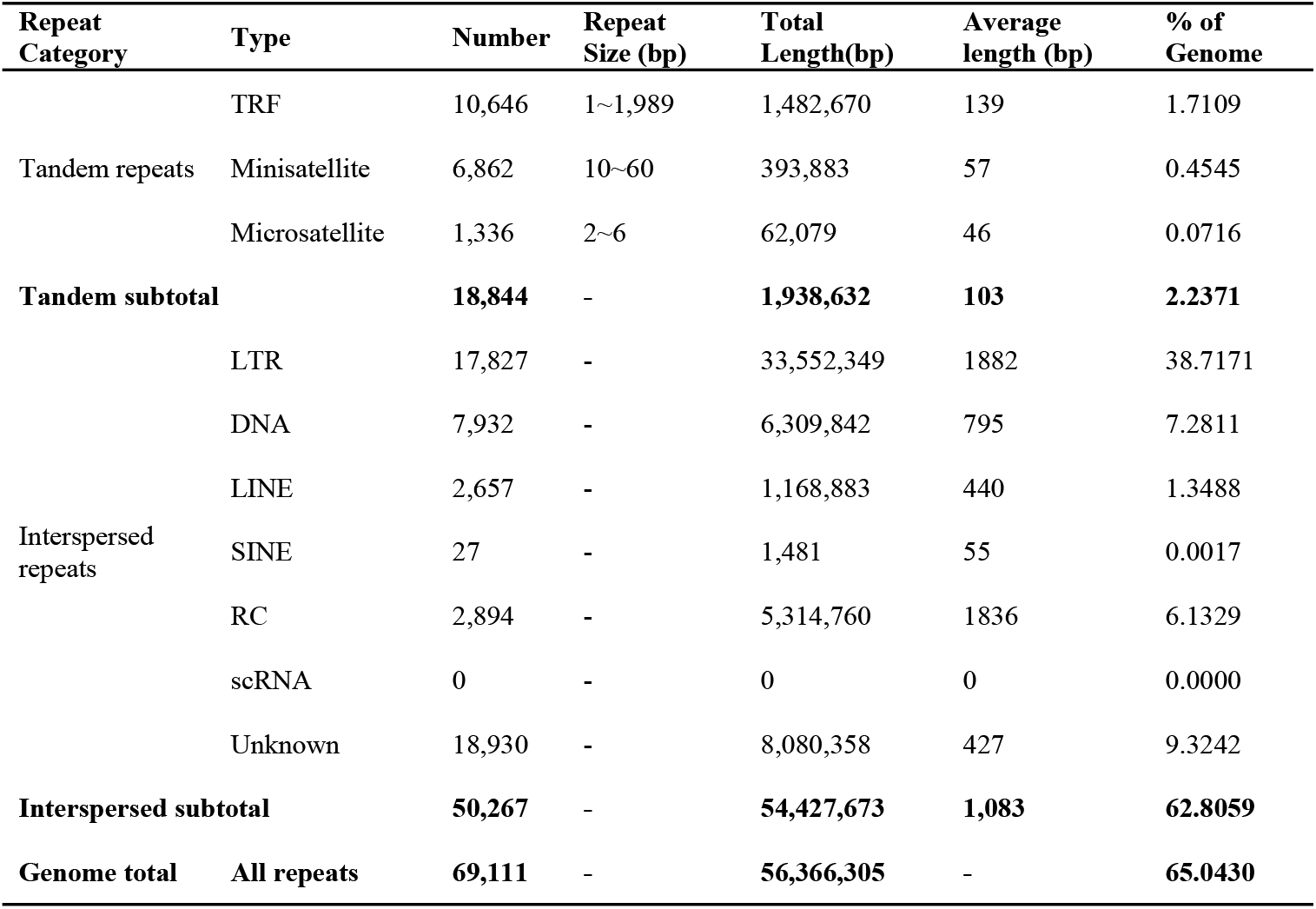
Repeats statistics in *Gp. lemaneiformis* genome.

When compared with closely related species, we noticed that 73% of the genome of *C. crispus* consists of repeated sequences, and the *Po. umbilicalis* had a substantial repeat element (43.9%) in its genome. Comparison of the repeat landscape of our genome and those in other red algae showed that the LTRs can be attributed to genome size variation.

### Gene predict and annotation

In total, we predicted 8,664 genes with an average length of 1.32 kb (Table 4). These protein-coding genes were further employed to analyze their functions using several public databases. We identified 8,202, 3,845 and 4,719 genes that showed homology to proteins in the NR, Swissport and COG databases, respectively. A total of 2,955 genes were assigned to GO classifications. Based on KEGG analysis, we annotated a total of 2,829 genes in the genome of *Gp. lemaneiformis*. Finally, 8,205 (94.70%) of the 8,664 predicted genes were annotated by at least one database.

**Table 4.**
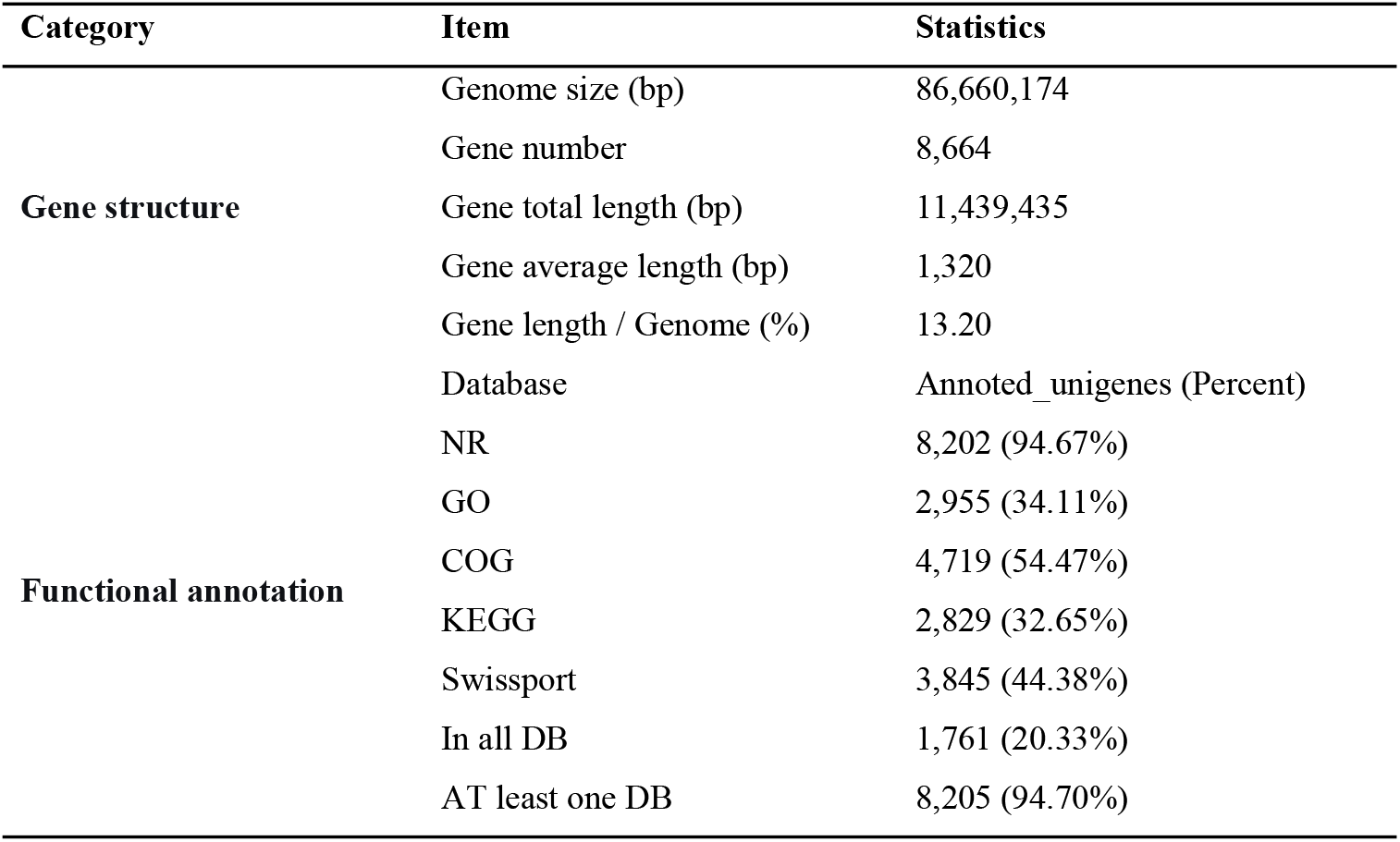
Predicted genes and function annotation statistics in *Gp. lemaneiformis* genome.

### Gene clusters and phylogenetic analysis

The protein sequences of *Gp. lemaneiformis* and 18 related species were compared to analyze the classification of species-specific genes between species. In *Physcomitrella patens*, 47,981 genes were grouped into 9,224 gene families, which was a greater number than in the other species (Table 5). The number of common families in all species was 510 (Figure 2B). The result in *Gp. lemaneiformis* exhibited 1,017 unique genes and 87 unique families (Table 5).

**Table 5.** Identification of gene family in multiple species.

| Species | Genes<br>number | Genes<br>in families | Unclustered<br>genes | Family<br>number | Unique<br>families | Common<br>families | Unique<br>genes |
| --- | --- | --- | --- | --- | --- | --- | --- |
| *Arabidopsis thaliana* | 20019 | 16851 | 3168 | 7195 | 1172 | 510 | 8553 |
| *Chlamydomonas reinhardtii* | 14483 | 11004 | 3479 | 8513 | 392 | 510 | 4790 |
| *Chlorella variabilis* | 9780 | 8021 | 1759 | 6332 | 190 | 510 | 2453 |
| *Chondrus crispus* | 9791 | 6697 | 3094 | 4928 | 259 | 510 | 4126 |
| *Coccomyxa subellipsoidea* | 9839 | 7774 | 2065 | 6275 | 212 | 510 | 2825 |
| *Cyanidioschyzon merolae* | 5014 | 4035 | 979 | 3682 | 37 | 510 | 1084 |
| *Cyanophora paradoxa* | 32108 | 14920 | 17188 | 6711 | 1801 | 510 | 25228 |
| *Ectocarpus siliculosus* | 24469 | 16764 | 7705 | 7902 | 1895 | 510 | 14264 |
| *Galdieria sulphuraria* | 7169 | 5909 | 1260 | 4266 | 233 | 510 | 2106 |
| *Gracilariopsis chorda* | 10806 | 9226 | 1580 | 7557 | 272 | 510 | 2500 |
| *Gracilariopsis lemaneiformis* | 9003 | 8257 | 746 | 7003 | 87 | 510 | 1017 |
| *Ostreococcus lucimarinus* | 7605 | 7020 | 585 | 6355 | 27 | 510 | 658 |
| *Ostreococcus tauri* | 7827 | 6400 | 1427 | 5961 | 50 | 510 | 1582 |
| *Phaeodactylum tricornutum* | 10408 | 8476 | 1932 | 6663 | 271 | 510 | 2842 |
| *Physcomitrella patens* | 47981 | 46759 | 1222 | 9224 | 1764 | 510 | 11053 |
| *Porphyra umbilicalis* | 13567 | 8118 | 5449 | 5278 | 487 | 510 | 7076 |
| *Pyropia yezoensis* | 7775 | 5654 | 2121 | 4066 | 262 | 510 | 3125 |
| *Thalassiosira pseudonana* | 11673 | 9183 | 2490 | 6889 | 371 | 510 | 3778 |
| *Volvox carteri* | 14426 | 11572 | 2854 | 8412 | 344 | 510 | 4641 |

**Figure 2.**
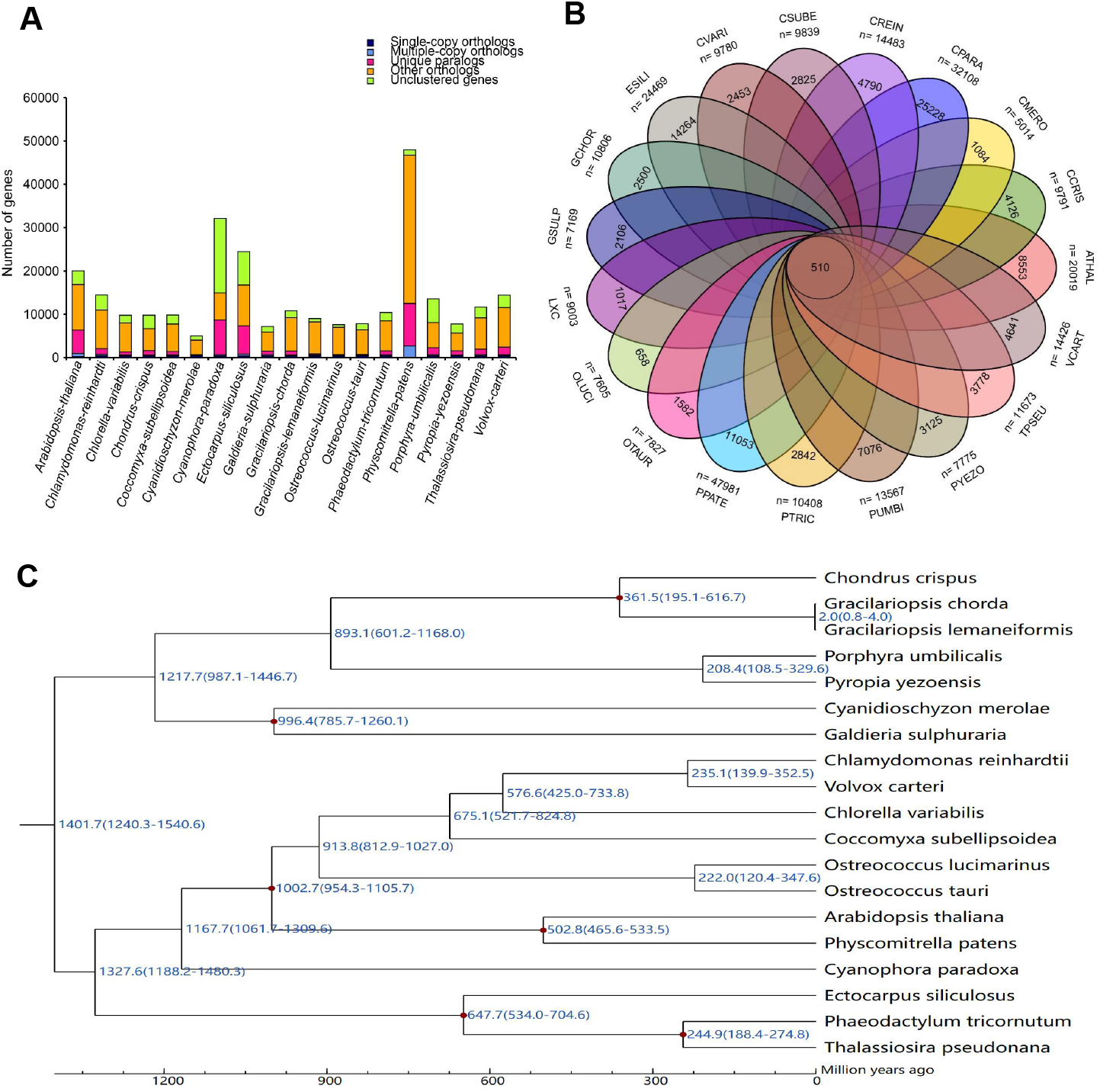
Comparative genomics and phylogeny of *Gp. lemaneiformis* across multiple species. (A)The distribution of orthologs and unique genes in multiple species. (B) A Venn diagram of common and unique gene families in multiple species. ATHAL, *Arabidopsis thaliana*; CREIN, *Chlamydomonas reinhardtii*; CVARI, *Chlorella variabilis*; CCRIS, *Chondrus crispus*; CSUBE, *Coccomyxa subellipsoidea*; CMERO, *Cyanidioschyzon merolae*; CPARA, *Cyanophora paradoxa*; ESILI, *Ectocarpus siliculosus*; GSULP, *Galdieria sulphuraria*; GCHOR, *Gracilariopsis chorda*; LXC, *Gracilariopsis lemaneiformis*; OLUCI, *Ostreococcus lucimarinus*; OTAUR, *Ostreococcus tauri*; PTRIC, *Phaeodactylum tricornutum*; PPATE, *Physcomitrella patens*; PUMBI, *Porphyra umbilicalis*; PYEZO, *Pyropia yezoensis*; TPSEU, *Thalassiosira pseudonana*; VCART, *Volvox carteri*. (C) Phylogenetic analysis of *Gp. lemaneiformis* and 18 other species.

The phylogenetic tree using the method of maximum-likelihood indicated that *Gp. lemaneiformis* shows a close relationship with *Gp. chorda*. To further investigate the divergence times of these species, the RelTime model was used. Fossil records were downloaded from the TIMETREE website (http://www.timetree.org) and used to calibrate the results. The divergence time of *Gp. lemaneiformis* and *G. chorda* was set to 2 million years ago (Figure 2C).

### Gene family expansion and contraction

Gene family expansion and contraction were analyzed for *Gp. lemaneiformis* and 8 additional species, based on the identified orthologs and the phylogenetic tree. The 9 species used for analysis include: *Gp. Lemaneiformis, Gp. chorda, Chondrus crispus, Porphyra umbilicalis, Cyanidioschyzon merolae, Galdieria sulphuraria, Ostreococcus lucimarinus, Chlamydomonas reinhardti* and *Arabidopsis thaliana*. Compared with the last common ancestor of *Gp. lemaneiformis* and *Gp. chorda*, 213 gene families were expanded and 327 gene families were contracted in *Gp. Lemaneiformis* (Figure 3A). Among the contracted families, 293 were completely lost, and 34 were retained but with reduced gene copies.

**Figure 3.**
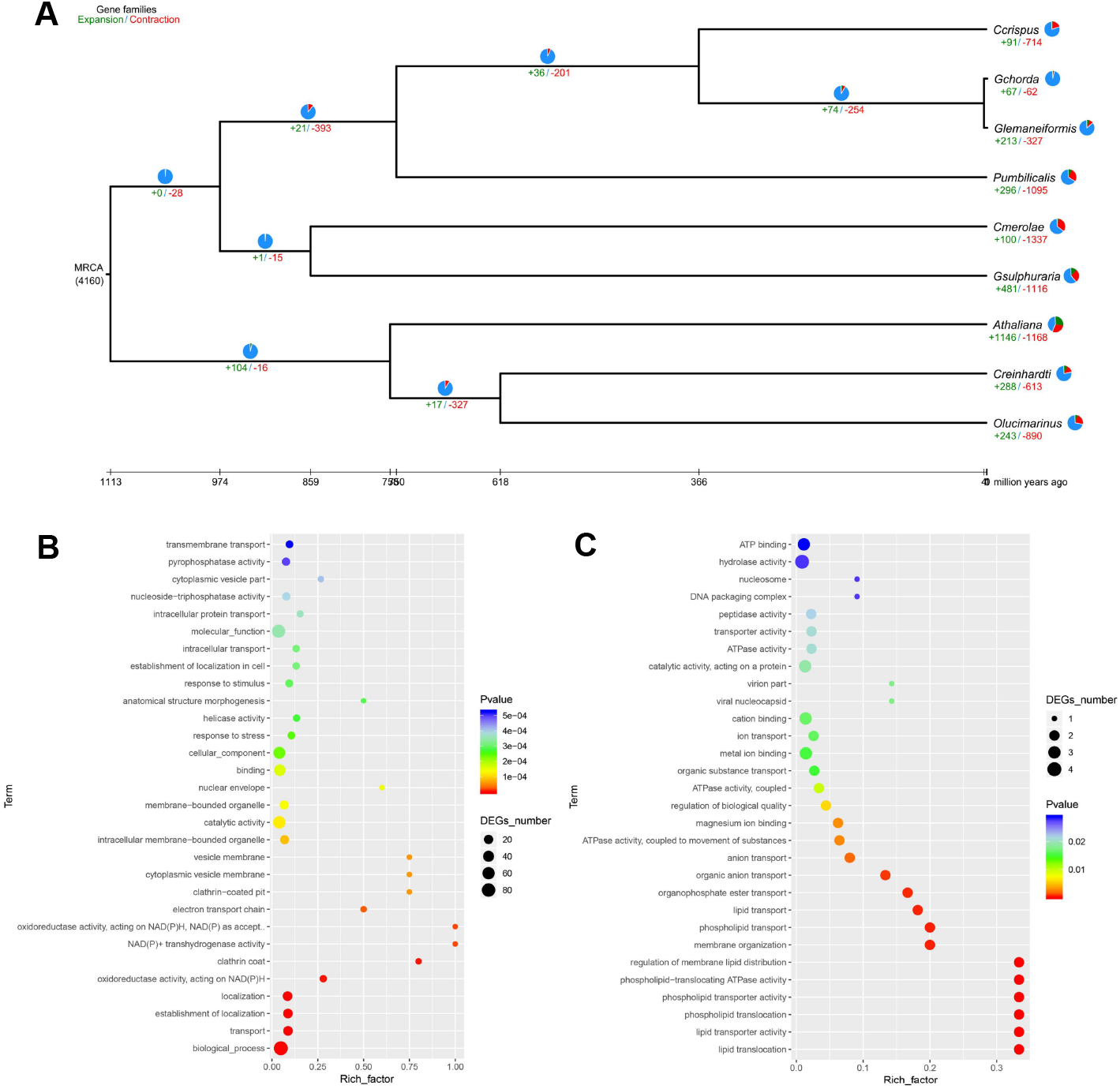
Expansion/contraction of gene families and their functional enrichment in *Gp. lemaneiformis*. (A) Phylogenetic analyses to reveal gene family expansion and contraction in multiple species. (B) GO terms enriched for expansions of gene families in *Gp. lemaneiformis*. (C) GO terms enriched for contraction of gene families in *Gp. Lemaneiformis*

Significantly expanded gene families were mainly enriched in NAD(P)H oxidoreductase activity, cytoplasmic vesicle membrane, stress response, and helicase activity (Figure 3B). Significantly contracted gene families were primarily enriched in lipid transporter activity, phospholipid transporter activity, phospholipid−translocating ATPase activity, and ion transport (Figure 3C).

Although *Gp. lemaneiformis* and *Gp. chorda* belong to the same genus *Gracilariopsis*, a large number of expanded and contracted gene families still exist between them. This suggests that after their divergence, different environmental pressures have shaped specific genetic variations to adapt to their respective habitats.

## Data Records

The raw data of Illumina, PacBio and Hi-C sequencing of *Gp. lemaneiformis* were deposited in the National Center for Biotechnology Information (NCBI) Sequence Read Archive (SRA) database with accession numbers SRR27204065, SRR20338037, SRR27638985. The transcriptomic sequencing data were deposited in the SRA at NCBI SRR27235189, SRR27235190, SRR27235188.

## Acknowledgements

This work was supported by the National Natural Science Foundation of China (32503178); the Shandong Provincial Natural Science Foundation (ZR2022QC105); the Fundamental Research Funds for the Central Universities (202313042); the China Agriculture Research System of MOF and MARA (CARS-50).

## Competing interests

The authors declare no competing interests.

## Notes

### Competing Interest Statement

The authors have declared no competing interest.

## References

1. Porse, H. & Rudolph, B. The seaweed hydrocolloid industry: 2016 updates, requirements,and outlook. J. Appl. Phycol. 29, 2187–2200 (2017).

2. Luo, H. T. et al. Protection of dietary selenium-enriched seaweed Gracilaria lemaneiformis against cadmium toxicity to abalone Haliotis discus hannai. Ecotoxicol. Environ. Saf. 171, 398–405 (2019).

3. Wei, Z. L. et al. A field scale evaluation of Gracilaria lemaneiformis co-cultured with Crassostrea gigas as a nutrient bioextraction strategy in Yantian Bay, China. Algal Res. 38, 101407 (2019).

4. Zhou, W. et al. Elevated-CO2 and nutrient limitation synergistically reduce the growth and photosynthetic performances of a commercial macroalga Gracilariopsis lemaneiformis. Aquaculture 550, 737878 (2022).

5. Wang, D. & Wu, F. X. China Fishery Statistical Yearbook (China Agricultural Press, 2024).

6. Kain, J. M. & Destombe, C. A review of the life history, reproduction and phenology of Gracilaria. J. Appl. Phycol. 7, 269–281 (1995).

7. Plastino, E. M. & de Oliveira F E.C., Deviations in the life-history of Gracilaria sp. (Rhodophyta, Gigartinales), from Coquimbo, Chile, under different culture conditions. Hydrobiologia 164, 67–74 (1988).

8. Liu, Y. T. et al. Cloning and transcription analysis of six members of the calmodulin family in Gracilaria lemaneiformis under heat shock. J. Appl. Phycol. 28, 643–651 (2016).

9. Liu, Y. T., Sun, H. Y., Ding, Y., Zang, X. N. & Zhang, X. C. A novel heat shock protein from Gracilariopsis lemaneiformis: gene cloning and transcription analysis in response to heat stress. J. Appl. Phycol. 30, 3623–3631 (2018).

10. Hu, Y. Y., Du, Q. W., Mi, P., Shang, E. L. & Sui, Z. H. Gene cloning and expression regulation in the pathway of agar and floridean starch synthesis of Gracilariopsis lemaneiformis (Rhodophyta). J. Appl. Phycol. 31, 1889–1896 (2019).

11. Wu, Q., Yin, J. R., Jiang, M., Zhang, J. Y. & Sui, Z. H. Identification, characterization and expression profiles of E2 and E3 gene superfamilies during the development of tetrasporophytes in Gracilariopsis lemaneiformis (Rhodophyta). BMC Genomics 24, 549 (2023).

12. Zhang, X. C. & van der Meer, J. P. A genetic study on Gracilaria sjoestedtii. Can. J. Bot. 66, 2022–2026 (2011).

13. Fu, F. et al. UV-irradiation mutation of tetraspores of Gracilariopsis lemaneiformis and screening of thermotolerant strains. J. Appl. Phycol. 26, 647–656 (2014).

14. Xiao, B. H., Hu, Y. Y., Feng, X. Q. & Sui, Z. H. Breeding of new strains of Gracilariopsis lemaneiformis with high agar content by ARTP mutagenesis and high osmotic pressure screening. Mar. Biotechnol. 25, 100–108 (2023).

15. Zhang, J. Y. et al. CRISPR/LbCas12a-mediated targeted mutation of Gracilariopsis lemaneiformis (Rhodophyta). Plant Biotechnol. J. 21, 235–237 (2023).

16. Wang, W. J., Wang, Z. Q., Lin, X. Z. & Xu, P. Characterization of Gracilaria lemaneiformis Bory (Gracilariaceae, Rhodophyta) cultivars in China using the total soluble proteins and RAPD analysis. Bot. Mar. 50, 177–184 (2007).

17. Pang, Q. Q., Sui, Z. H., Kang, K. H., Kong, F. N. & Zhang, X. C. Application of SSR and AFLP to the analysis of genetic diversity in Gracilariopsis lemaneiformis (Rhodophyta). J. Appl. Phycol. 22, 607–612 (2010).

18. Ding, H. Y., Sui, Z. H., Zhong, J., Zhou, W. & Wang, Z. X. Analysis and comparison on genetic diversity of haploid and diploid Gracilaria lemaneiformis populations from different places of Qingdao by AFLP. Period Ocean Univ. China 42, 99–105 (2012).

19. Hu, Y. Y. et al. Development of genomic simple sequence repeat markers and genetic diversity analysis of Gracilariopsis lemaneiformis (Rhodophyta). J. Appl. Phycol. 30, 707–716 (2018).

20. Hu, Y. Y. et al. Heterozygous single nucleotide polymorphic loci in haploid gametophyte of Gracilariopsis lemaneiformis (Rhodophyta). Front. Genet. 10, 1256 (2019).

21. Zhang, L. et al. Complete sequences of the mitochondrial DNA of the wild Gracilariopsis lemaneiformis and two mutagenic cultivated breeds (Gracilariaceae, Rhodophyta). PLoS One 7, e40241 (2012).

22. Du, Q. W., Bi, G. Q., Mao, Y. X. & Sui, Z. H. The complete chloroplast genome of Gracilariopsis lemaneiformis (Rhodophyta) gives new insight into the evolution of family Gracilariaceae. J. Phycol. 52, 441–450 (2016).

23. Zhou, W. et al. Genome survey sequencing and genetic background characterization of Gracilariopsis lemaneiformis (Rhodophyta) based on next-generation sequencing. PLoS One 8, e69909 (2013).

24. Sun, X. et al. Genomic analyses of unique carbohydrate and phytohormone metabolism in the macroalga Gracilariopsis lemaneiformis (Rhodophyta). BMC Plant Biol. 18, 94 (2018).

25. Matsuzaki, M. et al. Genome sequence of the ultrasmall unicellular red alga Cyanidioschyzon merolae 10D. Nature 428, 653–657 (2004).

26. Bhattacharya, D. et al. Genome of the red alga Porphyridium purpureum. Nat. Commun. 4, 1941 (2013).

27. Schönknecht, G. et al. Gene transfer from bacteria and archaea facilitated evolution of an extremophilic eukaryote. Science 339, 1207–1210 (2013).

28. Collén, J. et al. Genome structure and metabolic features in the red seaweed Chondrus crispus shed light on evolution of the Archaeplastida. Proc. Natl. Acad. Sci. 110, 5247–5252 (2013).

29. Brawley S. H. et al. Insights into the red algae and eukaryotic evolution from the genome of Porphyra umbilicalis (Bangiophyceae, Rhodophyta). Proc. Natl. Acad. Sci. 114, 6361–6370 (2017).

30. Ho, C. L., Lee, W. K. & Lim, E. L. Unraveling the nuclear and chloroplast genomes of an agar producing red macroalga, Gracilaria changii (Rhodophyta, Gracilariales). Genomics 110, 124–133 (2018).

31. Lee, J. et al. Analysis of the draft genome of the red seaweed Gracilariopsis chorda provides insights into genome size evolution in Rhodophyta. Mol. Biol. Evol. 35, 1869–1886 (2018).

32. Cao, M. et al. A chromosome-level genome assembly of Pyropia haitanensis (Bangiales, Rhodophyta). Mol. Ecol. Resour. 20, 216–227 (2020).

33. Wang, D. M. et al. Pyropia yezoensis genome reveals diverse mechanisms of carbon acquisition in the intertidal environment. Nat. Commun. 11, 4028 (2020).

34. Pflugmacher, S. & Steinberg, C. Activity of phase I and phase II detoxification enzymes in aquatic macrophytes. Angew. Bot. 71, 144–146 (1997).

35. Bolger, A. M., Lohse, M. & Usadel, B. Trimmomatic: a fexible trimmer for Illumina sequence data. Bioinformatics 30, 2114–2120 (2014).

36. Koren, S. et al. Canu: scalable and accurate long-read assembly via adaptive k-mer weighting and repeat separation. Genome Res. 27, 722–736 (2017).

37. Walker, B. J. et al. Pilon: an integrated tool for comprehensive microbial variant detection and genome assembly improvement. PLoS One 9, e112963 (2014).

38. Zhang, X. T., Zhang, S. C., Zhao, Q., Ming, R. & Tang, H. B. Assembly of allele-aware, chromosomal-scale autopolyploid genomes based on Hi-C data. Nat. Plants 5, 833–845 (2019).

39. Li, H. & Durbin, R. Fast and accurate long-read alignment with Burrows-Wheeler transform. Bioinformatics 26, 589–595 (2010).

40. Burton, J. N. et al. Chromosome-scale scaffolding of de novo genome assemblies based on chromatin interactions. Nat. Biotechnol. 31, 1119–1125 (2013).

41. Flynn, J. M. et al. RepeatModeler2 for automated genomic discovery of transposable element families. Proc. Natl. Acad. Sci. 117, 9451–9457 (2020).

42. Bao, Z. & Eddy, S. R. Automated de novo identifcation of repeat sequence families in sequenced genomes. Genome Res. 12, 1269–1276 (2002).

43. Price, A. L., Jones, N. C. & Pevzner, P. A. De novo identifcation of repeat families in large genomes. Bioinformatics 21, i351–i358 (2005).

44. Tempel, S. Using and understanding RepeatMasker. Methods Mol. Biol. 859, 29–51 (2012).

45. Benson, G. Tandem repeats fnder: a program to analyze DNA sequences. Nucleic Acids Res. 27, 573–580 (1999).

46. Haas, B. J. et al. Automated eukaryotic gene structure annotation using EVidenceModeler and the program to assemble spliced alignments. Genome Biol. 9, R7 (2008).

47. Stanke, M. & Morgenstern, B. AUGUSTUS: a web server for gene prediction in eukaryotes that allows user-defined constraints. Nucleic Acids Res. 33, W465–W467 (2005).

48. Birney, E., Clamp, M. & Durbin, R. GeneWise and Genomewise. Genome Res. 14, 988–995 (2004).

49. Kent, W. J. BLAT--the BLAST-like alignment tool. Genome Res. 12, 656–664 (2002).

50. Haas, B. J. et al. Improving the Arabidopsis genome annotation using maximal transcript alignment assemblies. Nucleic Acids Res. 31, 5654–5666 (2003).

51. Conesa, A. et al. Blast2GO: a universal tool for annotation, visualization and analysis in functional genomics research. Bioinformatics 21, 3674–3676 (2005).

52. Chen, F., Mackey, A. J., Stoeckert, C. J. & Roos, D. S. OrthoMCL-DB: querying a comprehensive multi-species collection of ortholog groups. Nucleic Acids Res. 34, 363–368 (2006).

53. Posada, D. JModelTest: phylogenetic model averaging. Mol. Biol. Evol. 25, 1253–1256 (2008) .

